# An open database of resting-state fMRI in awake rats

**DOI:** 10.1101/842807

**Authors:** Yikang Liu, Pablo D. Perez, Zilu Ma, Zhiwei Ma, David Dopfel, Samuel Cramer, Wenyu Tu, Nanyin Zhang

## Abstract

Rodent models are essential to translational research in health and disease. Investigation in rodent brain function and organization at the systems level using resting-state functional magnetic resonance imaging (rsfMRI) has become increasingly popular, owing to its high spatial resolution and whole-brain coverage. Due to this rapid progress, shared rodent rsfMRI databases can be of particular interest and importance to the scientific community, as inspired by human neuroscience and psychiatric research that are substantially facilitated by open human neuroimaging datasets. However, such databases in rats are still lacking. In this paper, we share an open rsfMRI database acquired in 90 rats with a well-established awake imaging paradigm that avoids anesthesia interference. Both raw and preprocessed data are made publically available. Procedures in data preprocessing to remove artefacts induced by the scanner, head motion, non-neural physiological noise are described in details. We also showcase inter-regional functional connectivity and functional networks calculated from the database.

## Introduction

Resting-state functional magnetic resonance imaging (rsfMRI) measures spontaneous brain activity in the absence of explicit external tasks or stimuli based on the blood-oxygenation-level dependent (BOLD) contrast (Biswal et al., 1995). Due to its high spatial resolution and whole-brain coverage, this method has tremendously advanced our understanding of human brain networks in terms of the function (Greicius et al., 2003; Hampson et al., 2002; Lowe et al., 1998), organization (Fox et al., 2005; Wang et al., 2010), developmental and aging profiles (Dosenbach et al., 2010; Fair et al., 2008), as well as genetic basis (Fu et al., 2015; Wiggins et al., 2012), and has provided a potential biomarker that can be used to track the progress of brain disorders, and evaluate the effects of treatment (Lee et al., 2013).

Relative to human research, studying rodent models using rsfMRI has unique advantages. First, environmental and genetic background are relatively uniform, making it relatively easier to separate their influences on brain networks and function (Gorges et al., 2017). Second, fMRI can be combined with cutting-edge neuroscience techniques such as electrophysiology (Majeed et al., 2011; Pan et al., 2010; Sloan et al., 2010), optogenetics (Desai et al., 2011; Lee et al., 2010; Liang et al., 2015b), calcium signal recording (Liang et al., 2017; Schlegel et al., 2018), Designer Receptors Exclusively Activated by Designer Drugs (DREADDs) (Grayson et al., 2016), which can facilitate bridging wide-range information from the cellular to systems levels. Applying rsfMRI in transgenic rodent models can further link imaging discoveries to neural mechanisms at genetic and molecular levels (Asaad and Lee, 2018). Third, rodent rsfMRI studies have high translational value. Using rsfMRI and other techniques, functional networks such as the thalamocortical and default mode networks have been identified in rodents that bear high anatomical resemblance as those in humans (Liang et al., 2013; Lu et al., 2012; Stafford et al., 2014). Topological organization such as small-worldness and rich-club organization is also well conserved in both humans and rodents (Bullmore and Sporns, 2009; Liang et al., 2011; Ma et al., 2018; van den Heuvel and Sporns, 2011). Taken together, rsfMRI provides a powerful tool in characterizing rodent models that complement human studies.

Despite these significant potentials, there is a large disparity in the number of publications between animal and human studies using rsfMRI. A major challenge is that anesthesia is often used in animal rsfMRI experiments to immobilize animals. It becomes increasingly clear that anesthesia changes physiological conditions (Tsukamoto et al., 2015), neurovascular coupling (Devor et al., 2007), brain metabolism (Hyder et al., 2002), and function of brain circuits and networks (Hamilton et al., 2017; Liang et al., 2011; Lu et al., 2007; Ma et al., 2017). In addition, the effects of anesthesia vary across different anesthetic agents and dosages (Grandjean et al., 2014; Hamilton et al., 2017), making it difficult to integrate data from different labs using different anesthetia protocols. Therefore, to avoid these issues it is important to image animals at the awake state when studying brain function.

Our lab has established an awake animal imaging paradigm that allows the brain function to be studied without the interference of anesthesia (Zhang et al., 2010). In this paradigm, animals were acclimated to the MRI scanning environment to minimize their stress and motion before imaging (King et al., 2005). We have demonstrated that this acclimation procedure, unlike studies of chronic stress that use prolonged daily restraint, does not induce chronic stress, nor does it interact with other stressors (Dopfel et al., 2019; Liang et al., 2014). By utilizing this method, we have investigated spatiotemporal dynamics of individual neural circuits (Liang et al., 2012a, 2013) and whole-brain networks (Liang et al., 2011; Liu and Zhang, 2019; Ma and Zhang, 2018; Ma et al., 2018). This method has also be employed to reveal changes in whole-brain connectivity architecture during brain development (Ma et al., 2018), under anesthesia (Hamilton et al., 2017; Liang et al., 2015a, 2012b; Ma et al., 2017; Smith et al., 2017), as well as neuroplastic changes induced by traumatic stress (Dopfel et al., 2019; Liang et al., 2014) and drugs (Crenshaw et al., 2015; Pérez et al., 2018; Roses et al., 2014). Taken together, these studies have demonstrated the validity and value of the awake fMRI approach.

In this paper, we share an open database to the public, which contains 168 rsfMRI scans from 90 rats acquired in the awake state. We provide both raw and preprocessed data. Some results obtained from routine analyses are demonstrated.

## Methods and Materials

### Animals

Data were acquired in 90 adult male Long-Evans rats (300 g–500 g), part of which were used in previous publications (Dopfel et al., 2019; Liu and Zhang, 2019; Ma et al., 2018; Ma and Zhang, 2018; Ma et al., 2018). All rats were housed in Plexiglas cages (two per cage) with food and water provided *ad libitum*. A 12 h light: 12 h dark schedule and temperature between 22 °C and 24 °C were maintained. All experiments were approved by the Institutional Animal Care and Use Committee (IACUC) of the Pennsylvania State University.

### Acclimation procedure

The purpose of this procedure is to acclimate the animal to the restraining system as well as the noisy and confined environment inside the MRI scanner. Details of the acclimation procedure can be found in our previous publications (Dopfel and Zhang, 2018; Gao et al., 2017). Briefly, EMLA cream (2.5% lidocaine and 2.5% prilocaine) was applied topically to the nose, jaw, and ear areas to relieve any discomfort associated with the restrainer 20 min prior to the procedure. The animal was then briefly anesthetized with 2-4% isoflurane and placed in a head restrainer, in which the teeth and nose were secured by a bite bar and a nose bar, respectively, and ears were secured by adjustable ear pads. Forepaws and hindpaws were loosely taped to prevent the animal from accidental self-injury. After that, the body was placed in a Plexiglas body holder with the shoulders secured by a pair of shoulder bars. The whole system allowed unrestricted respiration. Isoflurane was discontinued after the setup. The restrainer was then fixed to a body holder. After the animal woke up, the system was placed into a black opaque chamber where the prerecorded sound from various imaging sessions was played. The animal was acclimated for 7 days with an incremental exposure time up to 60 min (i.e. 15, 30, 45, 60, 60, 60 and 60 min from Day 1 to Day 7, respectively).

### Data acquisition

Data were acquired on a 7T Bruker 70/30 BioSpec running ParaVision 6.0.1 (Bruker, Billerica, MA) at the High Field MRI Facility at the Pennsylvania State University. Similar to the acclimation procedure, the animal was briefly anesthetized with 2-4% isoflurane and were placed in a head restrainer integrated with a birdcage head coil. The isoflurane was discontinued once the setup was finished. rsfMRI acquisition started when the animal was fully conscious (usually within 10-15 min). A single-shot gradient-echo echo-planar imaging (GE-EPI) sequence was used with the following parameters: repetition time (TR) = 1000 ms; echo time (TE) = 15 ms; matrix size = 64 × 64; field of view (FOV) = 3.2 × 3.2 cm^2^; slice number = 20; flip angle = 60°, 600, 900, or 1200 volumes per scan. A representative raw EPI image was shown in Fig. S1. Anatomical images were also acquired with a rapid imaging with refocused echoes (RARE) sequence with the following parameters: TR = 1500 ms; TE = 8 ms; matrix size = 256 × 256; FOV = 3.2 × 3.2 cm^2^; slice number = 20.

### Data preprocessing

The preprocessing pipeline is outlined in Fig. 1, which included 7 steps:

1. Volumes with excessive motion were discarded (i.e. scrubbing).
2. rsfMRI images were manually co-registered to an anatomical template with rigid-body transformation.
3. Co-registered images were corrected for head motion and motion parameters were recorded.
4. Non-neural artefacts were identified with independent component analysis (ICA). Time courses of noise independent components (ICs) were recorded.
5. The motion-corrected images were spatially smoothed.
6. The spatially-smoothed images were softly cleaned by regressing out noise IC time courses, motion parameters and the signals from the white matter (WM) and cerebral spinal fluid (CSF).
7. The soft-cleaned images were temporally bandpass filtered.

**Figure 1.**
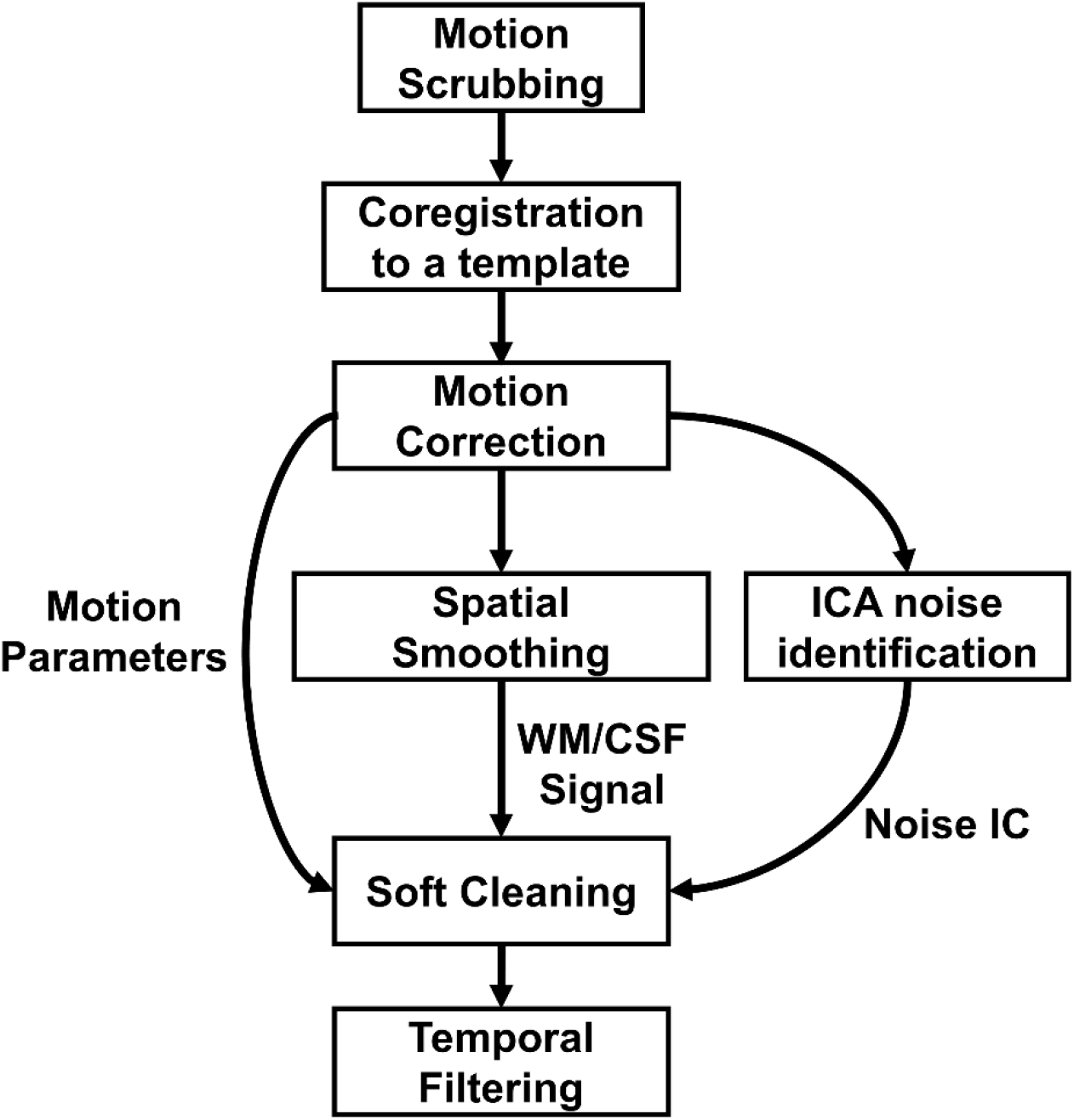
rsfMRI data preprocessing pipeline.

All the source codes used for preprocessing can be downloaded from the GitHub repository: https://github.com/liu-yikang/rat_rsfmri_preprocessing. Details of each step were described below.

### Motion scrubbing and co-registration

First, motion was evaluated by calculating the relative framewise displacement (FD) of each rsfMRI volume (Power et al., 2012). Specifically, the geometric transformation from each frame (i.e. 3D volume) to the first frame were evaluated by the built-in function *imregtform* in MATLAB (The Mathworks Inc., Natick, MA, USA) with six degrees of transformation considered (i.e. rigid-body transformation), including translations in the three orthogonal axes (translation distances for the frame *i* are denoted as *x_i_*, *y_i_*, and *z_i_*) and rotations around the three axes (rotation angles are denoted as *α_i_*, *β_i_*, and *γ_i_*). Then *FD_i_* = | Δ*x_i_* | + | Δ*y_i_* | + | Δ*z_i_* | + *r* · (| Δ*α_i_* | + | Δ*β_i_* | + | Δ*γ_i_* |), where *r* = 5 mm, which is approximately the distance measured from the cortex to the center of the rat head. Frames with *FD* > 0.2 mm and their neighbor frames were discarded. The first 10 frames of each scan were also discarded to ensure steady state of magnetization. Scans with less than 90% of the total number of frames left were excluded from further analysis. This procedure and parameters used can effectively minimize motion artefacts as confirmed in our previous studies (Liu and Zhang, 2019; Ma and Zhang, 2018).

Next, the first frame of each rsfMRI scan was manually co-registered (i.e. aligned) to a T2-weighted anatomical template using an in-house software written in MATLAB. To ensure the quality of alignment, voxels at brain boundaries, ventricles, and WM in the anatomical template were displayed as landmarks on a graphical-user interface (Fig. S2). After the coregistration of the first frame, the same geometric transformation was applied to the remaining frames.

Subsequently, head motions were corrected using SPM12 (http://www.fil.ion.ucl.ac.uk/spm/) with a dilated brain mask applied, in which each frame was co-registered to the first frame through a rigid-body transformation. Motion parameters were recorded for further use. The distribution of averaged FD across scans is displayed in Fig. 2

**Figure 2.**
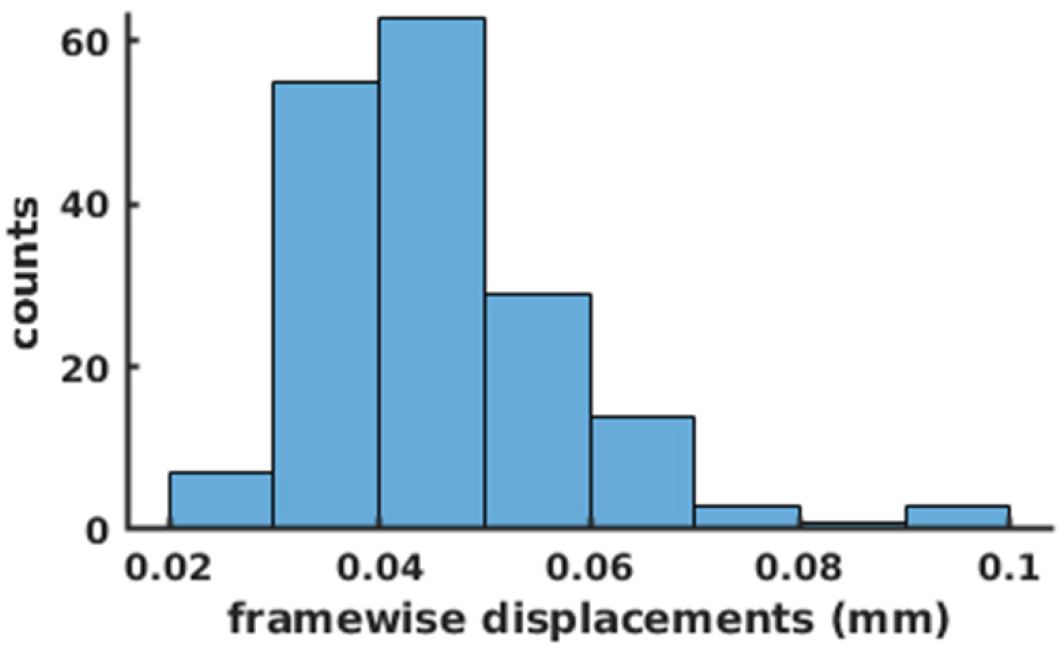
Distribution of averaged frame-wise displacement across scans.

### ICA-based artefact identification

We used ICA to remove non-neural artefacts potentially related to motion, breathing, heartbeats, and/or scanner instability from spatially co-registered data. ICA-based artefact removal has been widely applied in human and rodent studies (Griffanti et al., 2015, 2014; Han et al., 2019; Salimi-Khorshidi et al., 2014; Smith et al., 2013). It leverages the independency between spatial and/or temporal patterns of the neural and non-neural components to separate them. In this method, users can manually identify each IC as real signal or noise based on their spatial, temporal, and spectral features. Standards of manually classifying ICs with these features for the HCP data (Smith et al., 2013) were listed in Table 1 (Griffanti et al., 2017).

**Table 1.**
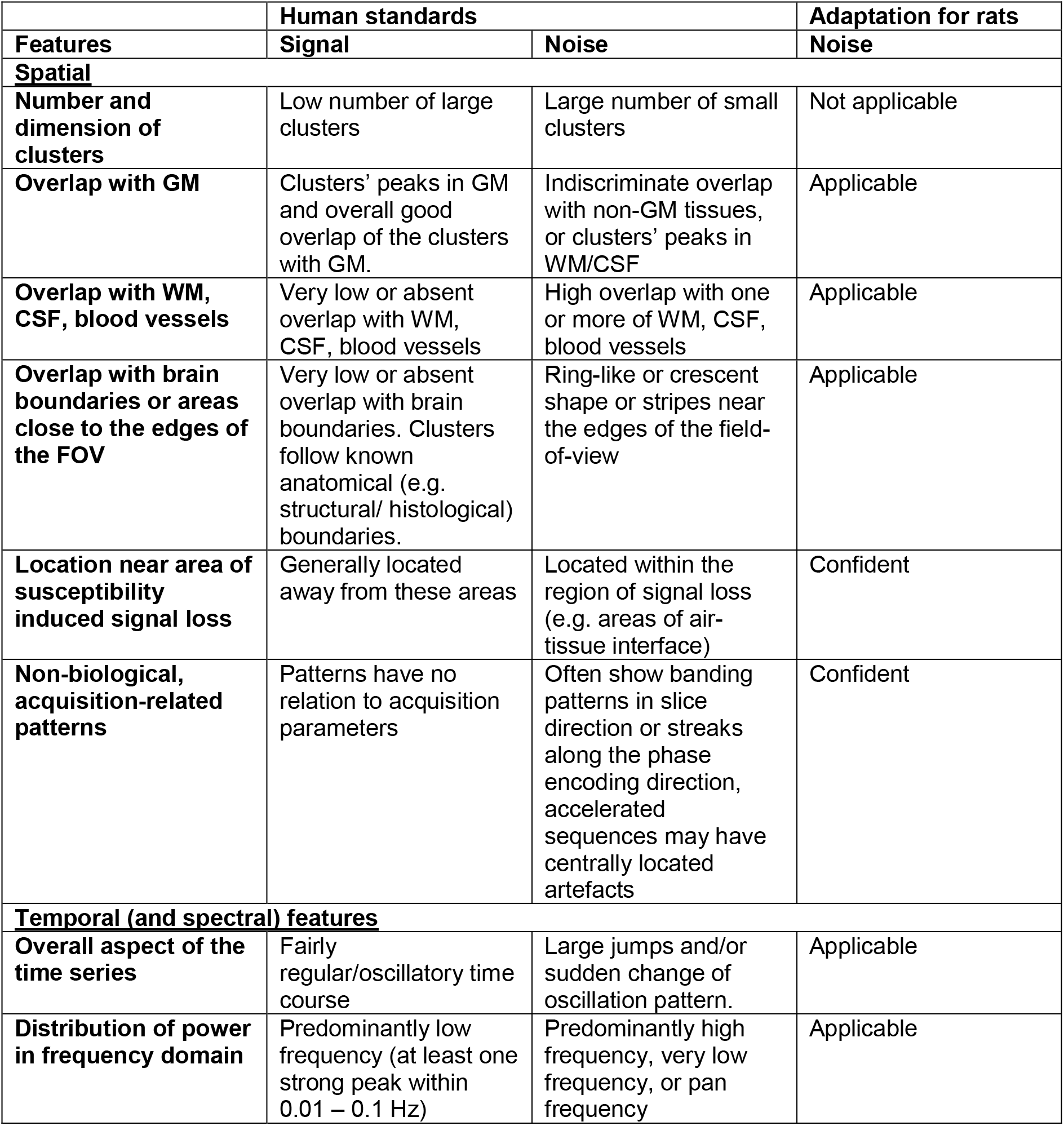
Features of signal- and noise-related independent components.

|  | **Human standards** |  | **Adaptation for rats** |
| --- | --- | --- | --- |
| **Features** | **Signal** | **Noise** | **Noise** |
| **<u>Spatial</u>** |  |  |  |
| **Number and dimension of clusters** | Low number of large clusters | Large number of small clusters | Not applicable |
| **Overlap with GM** | Clusters’ peaks in GM and overall good overlap of the clusters with GM. | Indiscriminate overlap with non-GM tissues, or clusters’ peaks in WM/CSF | Applicable |
| **Overlap with WM, CSF, blood vessels** | Very low or absent overlap with WM, CSF, blood vessels | High overlap with one or more of WM, CSF, blood vessels | Applicable |
| **Overlap with brain boundaries or areas close to the edges of the FOV** | Very low or absent overlap with brain boundaries. Clusters follow known anatomical (e.g. structural/ histological) boundaries. | Ring-like or crescent shape or stripes near the edges of the field-of-view | Applicable |
| **Location near area of susceptibility induced signal loss** | Generally located away from these areas | Located within the region of signal loss (e.g. areas of air-tissue interface) | Confident |
| **Non-biological, acquisition-related patterns** | Patterns have no relation to acquisition parameters | Often show banding patterns in slice direction or streaks along the phase encoding direction, accelerated sequences may have centrally located artefacts | Confident |
| **<u>Temporal (and spectral) features</u>** |  |  |  |
| **Overall aspect of the time series** | Fairly regular/oscillatory time course | Large jumps and/or sudden change of oscillation pattern. | Applicable |
| **Distribution of power in frequency domain** | Predominantly low frequency (at least one strong peak within 0.01 – 0.1 Hz) | Predominantly high frequency, very low frequency, or pan frequency | Applicable |

Our specific procedure followed these guidelines, but was also adapted to the characteristics of rat data. We found one criterion not applicable to our rat data. In the HCP guideline, signal ICs tend to have a few large clusters, whereas noise ICs tend to have many small clusters. This did not always hold true in rat data due to the relative portion of the cortex versus sub-cortex. The human brain is dominantly composed of the cortex, which contributes to most clustered structures in signal ICs. In contrast, 2/3 volume of the rat brain is sub-cortex that includes numerous heterogeneous nuclei. Thus, neural components in rats may not always display large clusters. Therefore, we grouped the HCP criteria into three categories: not applicable; applicable; confident, as listed in Table 1 and used the following criterion to label noise/signal ICs: an IC was classified as a noise component if it had one or two “confident” features or had at least two of the following three “applicable” features: 1) its spatial map is located predominately at white matters, ventricles, or brain boundaries; 2) its time course has sudden jumps and/or change of fluctuation pattern; 3) the frequency spectrum is flat or dominated by very low or high frequency.

Prior to ICA, we spatially smoothed each frame with a Gaussian kernel with full width at half maximum (FWHM) = 0.7 mm. The size of the kernel was determined empirically to reduce noise but still maintain the difference between neural and non-neural components. After that, spatial ICA was separately conducted on each scan using the GIFT ICA toolbox (Calhoun et al., 2009) with the number of ICs set at 50. Subsequently, we calculated the time courses of ICs by regressing their spatial maps against each frame.

We manually labeled ICs as signal or noise components for all scans of all animals based on the features of their spatial maps (z-scored, thresholded at z > 2), time courses, and spectra using the criterion mentioned above (Fig. 3). Two representative noise ICs were demonstrated (Fig. 3).

**Figure 3.**
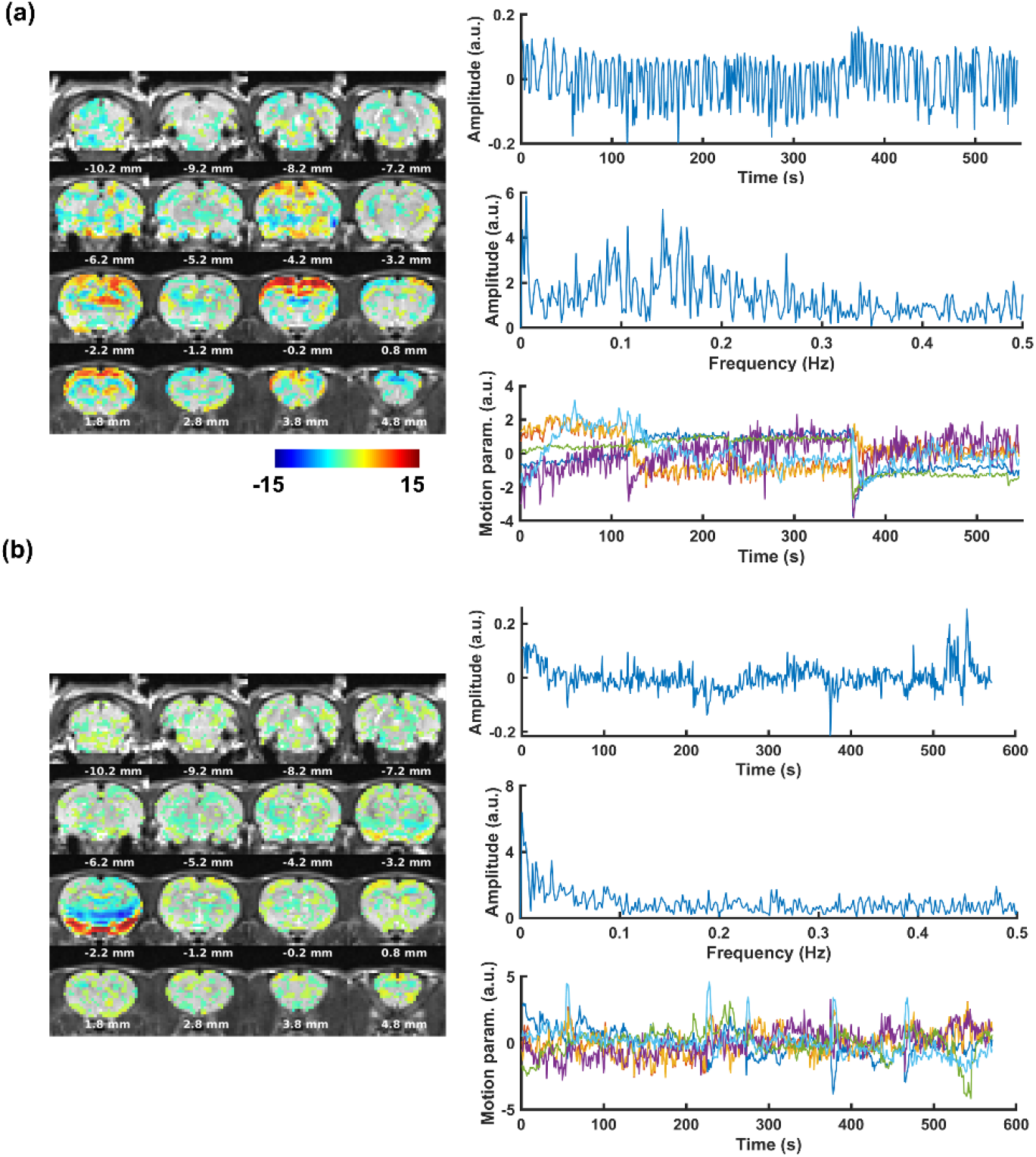
Two representative noise ICs from ICA-based cleaning.

### Spatial smoothing, soft cleaning, and temporal filtering

After identifying noise ICs, the original motion-corrected but unsmoothed rsfMRI data were spatially smoothed with a Gaussian kernel (FWHM=1 mm). WM and CSF signals were calculated by averaging the signal of voxels in the WM or ventricles, respectively.

We “softly” removed the noise ICs obtained using the method proposed by (Griffanti et al., 2014), and regressed out six motion parameters obtained in motion correction as well as the WM/CSF signals from the rsfMRI data. In this method, only the unique parts of variance explained by the noise ICs were removed, whereas parts shared with the signal ICs were reserved. Briefly, first all ICA time courses and rsfMRI images were regressed by the corresponding motion parameters and WM/CSF signals, resulting in regressed ICA time courses (*ICA_m_*) and regressed images (*Y_m_*). Second, we regressed *ICA_m_* against *Y_m_* to obtain the weight of unique contribution of each IC to the data: *β* = pinv(*ICA_m_*) · *Y_m_*. Third, we removed the unique contribution of the noise components from the data: *Y_clean_* = *Y_m_* − *ICA_m_*(noise) · *β*(noise).

Finally, the signal of each voxel in each rsfMRI scan was temporally filtered with a 4^th^-order bandpass Butterworth filter (0.01 – 0.1 Hz).

All raw and preprocessed data have been uploaded and can be freely downloaded from: https://nitrc.org/projects/rat_rsfmri. The folder structure of raw and preprocessed data is described in the Appendix.

## Results

### Region-based correlational analysis

Fig. 4a shows the group-level pairwise FC between 90 regions of interest (ROIs) covering the whole brain based on Swanson atlas (Swanson, 2004). ROIs were organized and color coded by the brain systems, indicated by the bars next to the FC matrix. The group-level FC (in *t* value) was calculated by fitting a linear mixed model (subject variability modeled as the random effect) to the FC of individual scans (i.e. one-sample *t* tests on the random effect), which was quantified as Fisher Z-transformed Pearson correlation coefficient of the averaged rsfMRI signals between every two ROIs. To ensure the same degree of freedom of individual scans, each scan with 600 or 900 volumes was truncated into a 540-volume scan, and each scan with 1200 volumes was truncated into two 540-volume scans. This operation resulted in 181 scans for processing. The lower triangle shows entries (i.e. connections) with significant FC (*p* < 0.05, false discovery rate (FDR) corrected). Fig. 4b shows the group-level reproducibility of FC, calculated by the similarity of FC matrices in two randomly divided subgroups. The correlation of the corresponding off-diagonal entries between the two matrices was 0.937 (p ≈ 0, Fig. 4c). The reproducibility of FC at the individual level was quantified by the correlation of off-diagonal entries between the FC matrix of each individual animal and that of the whole group (excluding the tested animal). The averaged correlation value across animals was 0.639 ± 0.090 (mean ± std, p ≈ 0).

**Figure 4.**
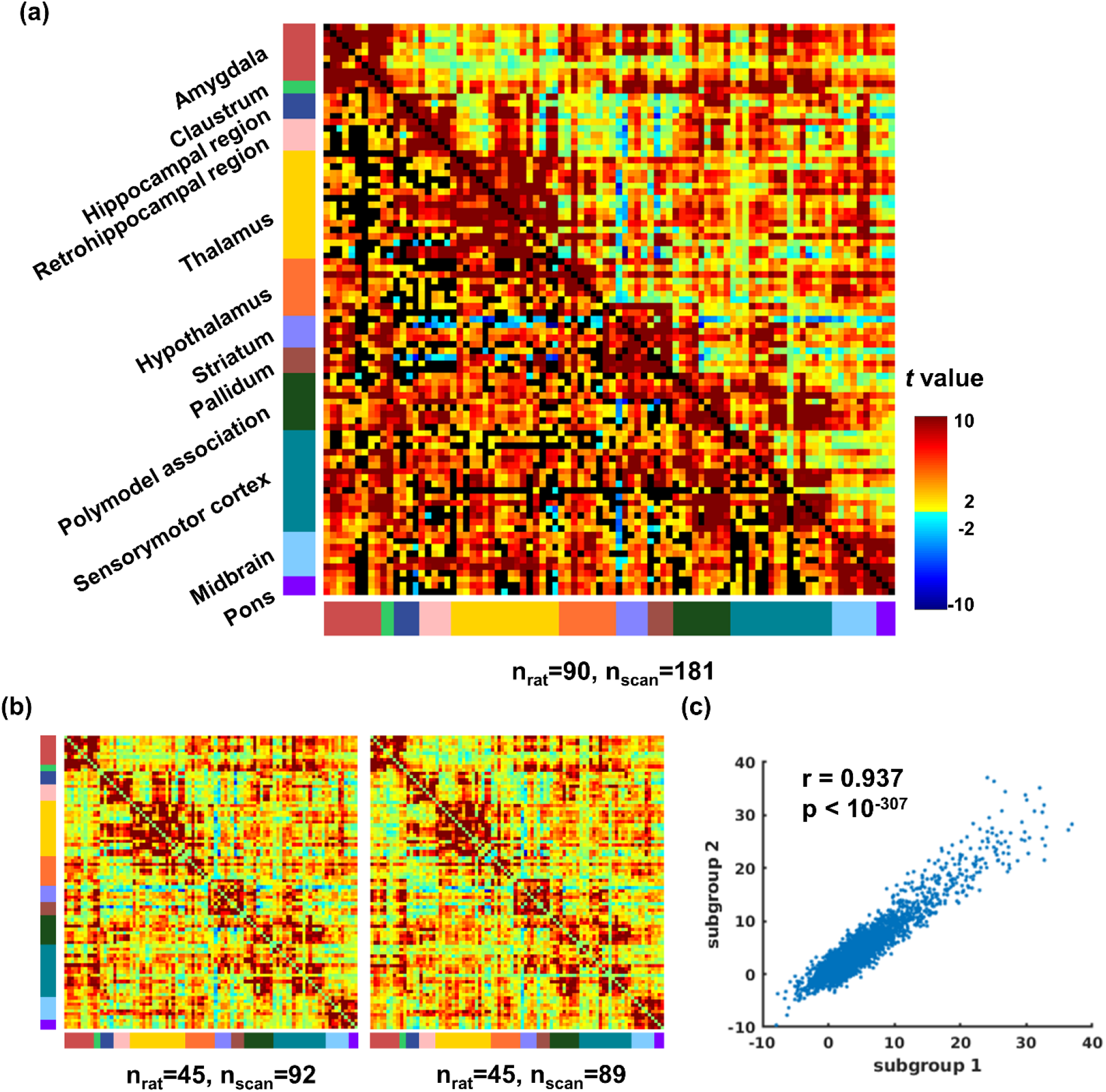
FC between ROIs defined by the Swanson atlas. (a) FC matrix of all animals. The lower triangle shows connections (i.e. entries) with significant FC thresholded at p < 0.05, FDR-corrected. (b) FC of two randomly divided non-overlapping subgroups. (c) Correlation of the corresponding off-diagonal entries in the two matrices in (b).

Fig. 5 shows a few examples of group-level seed maps with the seeds in the visual cortex (VIS), primary motor cortex (MOp), primary somatosensory cortex (SSp), retrosplenial cortex (RSP), insula cortex (INS), and infralimbic cortex (IL), respectively. Voxel-wise FC (in *t* value) was calculated in the same way. All other seed maps can be downloaded from the database.

**Figure 5.**
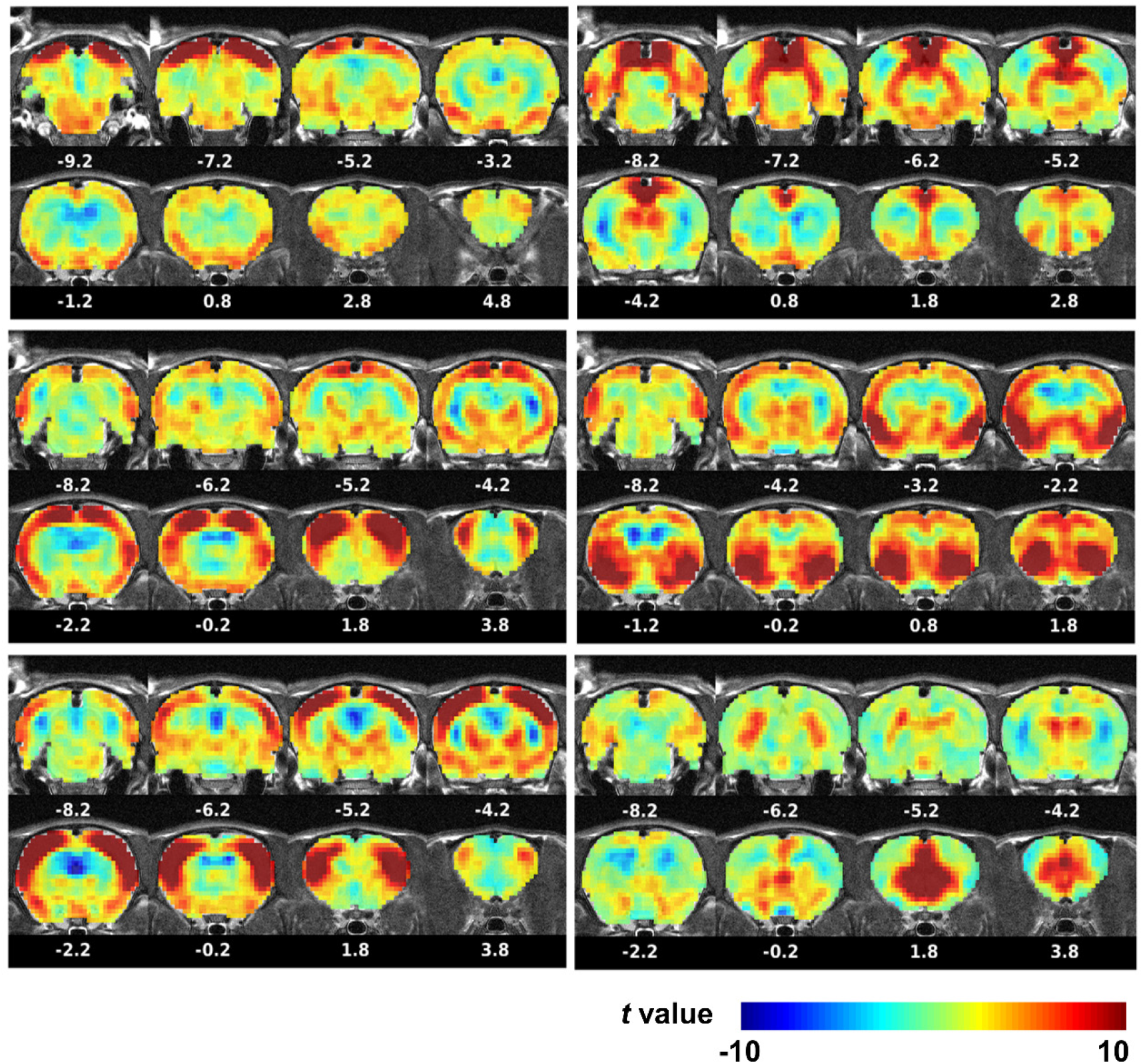
Representative seedmaps. By column, the seed regions are the visual cortex, primary motor cortex, and primary somatosensory cortex; retrosplenial cortex, insular cortex, and infralimbic cortex.

### ICA analysis

In this section, we demonstrate functional networks revealed by ICA. Using the GIFT toolbox, we ran spatial group ICA on all preprocessed rsfMRI scans with the number of components set at 75. Fig. 6 shows the spatial patterns of all ICA components, thresholded at z > 7 (p < 0.00001).

**Figure 6.**
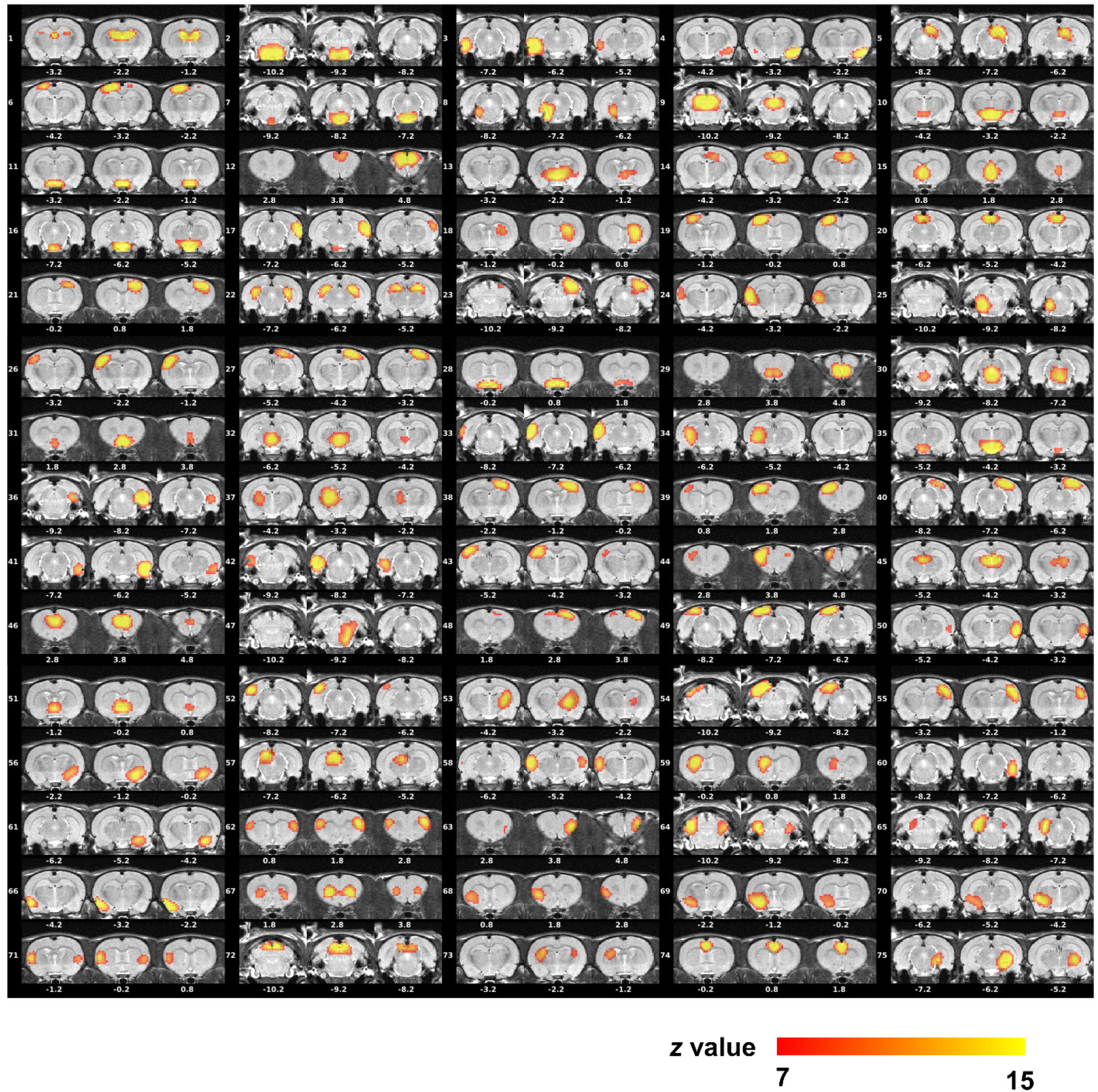
Spatial maps of 75 ICs generated by spatial group ICA. All maps are thresholded at z=7 (p<0.00001).

We also demonstrate the connectivity architecture between ICA components. The FC between every two ICA components was determined by the Pearson correlation between their time courses for each rsfMRI scan (truncated in the same manner as described above). Group-level inference was determined using the same linear mixed model, resulting in a *t* value for each pair of ICA components, displayed in Fig. 6a. The lower triangle of the *t* matrix shows significant entries (i.e. between-component connections) thresholded at p < 0.05, FDR-corrected. Subsequently, all components were hierarchically clustered with the Ward’s method (Murtagh and Legendre, 2014) using FC as the similarity. The dendrogram is shown in Fig. 7a (top), where we cut off the dendrogram with an empirical threshold, which resulted into 4 modules.

**Figure 7.**
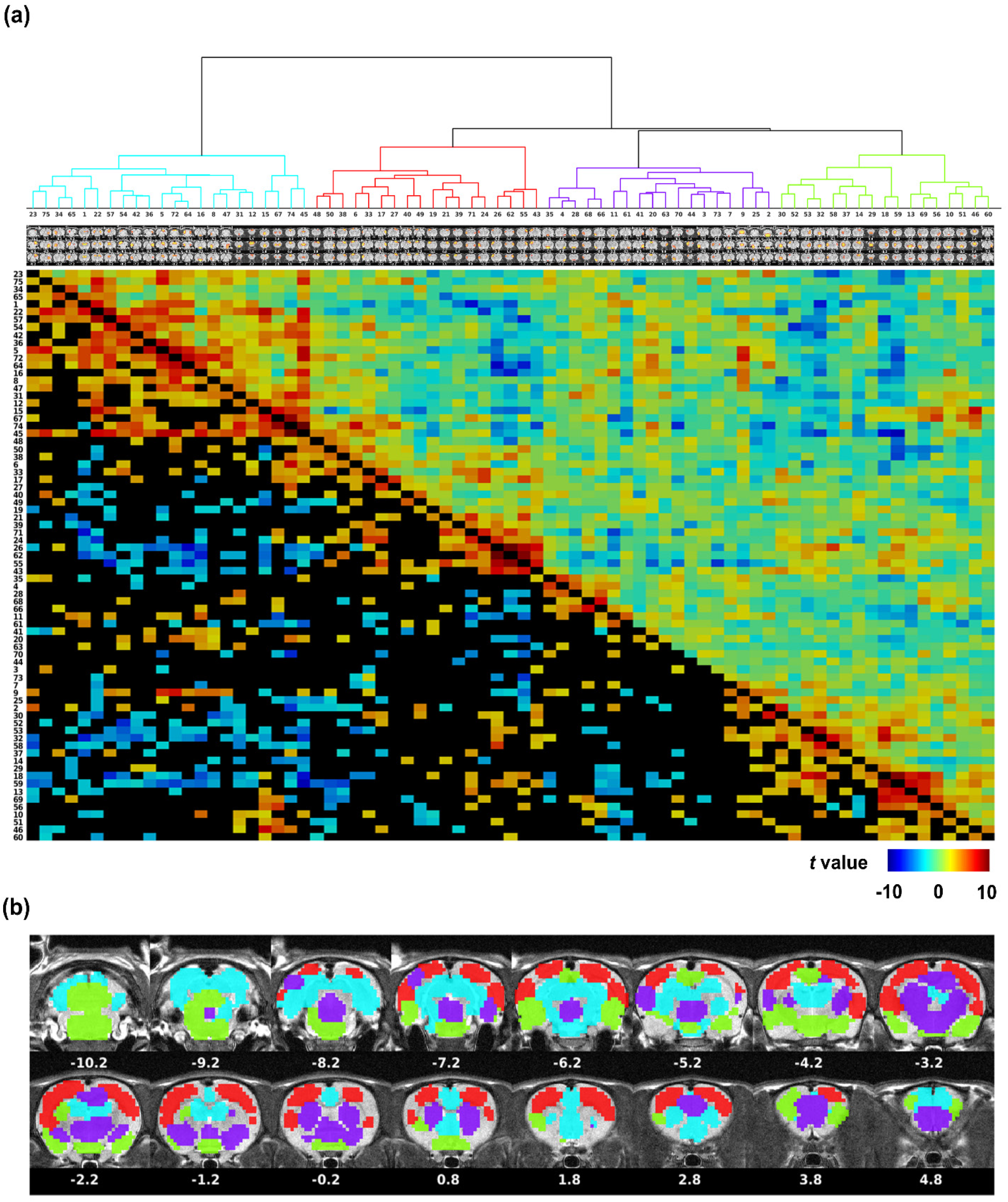
Connectional structure between all ICA components. (a) Hierarchical clustering of ICs (upper panel), and between-IC FC matrix (lower panel). The dendrogram was cut off with an empirical threshold, resulting into 4 modules. The lower triangle of the FC matrix was thresholded by the FDR-corrected p value of 0.05. (b) Community structures revealed by color coding ICs based on their corresponding communities (z>10).

An interesting observation is that one module (blue module, including the VIS, hippocampus (HC), superior colliculus (SC), inferior colliculus (IC), temporal association cortex (TeA), ectorinal cortex (ECT), paraflocculus (PFl), flocculus (Fl), pontine nuclei (Pn), dorsal thalamus (THA), olfactory cortex (OC), nucleus accumbens (ACB), anterior cingulate cortex (ACC), and orbital cortex (ORB)) exhibits pronounced anticorrelated FC with two other modules (red module, including the MO, SS, auditory cortex (AUD), and parietal cortex (PC), as well as the green module, including the Pn, VIS, dorsolateral THA, AUD, caudate putamen (CPu), piriform cortex (Pir), hypothalamus (HYPO), OC, paralimbic cortex (PL), and IL).

### Effects of the ICA cleaning

Finally, we demonstrate the effects of ICA cleaning. Fig. 8 shows the effects of regressing out noise ICs on FC measurement. We compared FC in ICA-cleaned and uncleaned data (without soft regression of noise components). Fig. 8a shows ROI-wise FC in uncleaned data. Fig. 8b shows the FC distributions with and without ICA cleaning, demonstrating that ICA cleaning increased the significance level of FC (i.e. increased t values).

**Figure 8.**
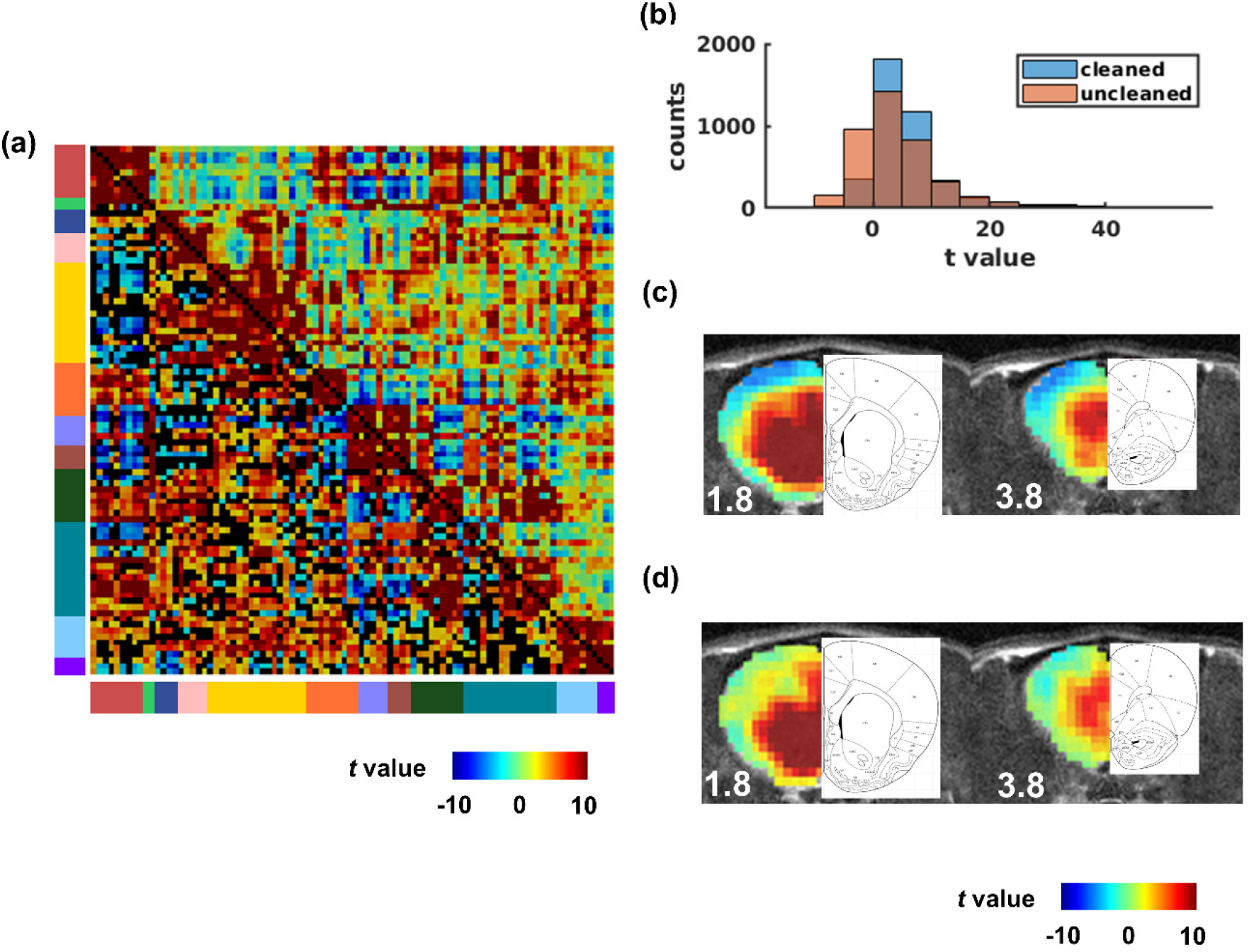
Effects of ICA cleaning. (a) ROI-wise FC in the uncleaned data. (b) Histogram of ROI-wise FC of cleaned and uncleaned data. (c) and (d) uncleaned and cleaned seedmaps of nucleus accumbens, respectively.

Figs. 8c and 8d show the ACB seedmaps without and with ICA cleaning, respectively. The anatomical definition on the same slices was juxtaposed for comparison. It is clear that ICA cleaning was able to remove artifacts at the top brain boundary. Regions showing prominent FC in both seedmaps were spatially more specific in relation to the anatomical structure after ICA cleaning.

## Discussion

Neuroscience and psychiatric research have been substantially facilitated by open neuroimaging datasets (Poldrack and Gorgolewski, 2017; Thompson et al., 2014; Van Essen et al., 2013). Data sharing not only speeds up scientific discoveries by leveraging a high statistical power brought by large volumes of data, but also incentivizes researchers to develop new analysis methods that can be tested on these datasets. While a large number of open databases of human rsfMRI studies have been established, such database in rodents, particularly awake rodents are lacking. Considering that rodent models are an important translational tool for clinical and basic neuroscience research, here we share an open rsfMRI database acquired in 90 awake rats in the neuroscience and neuroimaging communities, and illustrate the data acquisition protocol and preprocessing procedure.

There has been growing interest in studying brain function and organization in awake rodents using rsfMRI, which avoids interference of anesthesia and permits correlation to behavioral data (Bergmann et al., 2016; Brydges et al., 2013; Chang et al., 2016; Liang et al., 2011; Stenroos et al., 2018; Yoshida et al., 2016). One major challenge of awake rodent fMRI is to control motion and stress during data acquisition. We addressed the issue in three aspects: first, we used a 3D-printed head restraint system to limit animals’ head motion; second, we adopted a 7-day acclimation routine prior to imaging, which has been shown to significantly reduce stress during image acquisition (King et al., 2005); third, we used stringent data preprocessing including scrubbing volumes with excessive motion, regressing out motion parameters, WM/CSF signals, and non-neural artefacts using ICA cleaning.

Our data demonstrate high inter-subject reproducibility in whole-brain FC matrices both at the group level and the individual level. We showcase inter-regional functional connectivity and functional networks calculated from the database, including the motor, visual, somatosensory, and auditory networks. In our library we include seed maps from all individual anatomical ROIs. Taken together, the database shared should provide a resource for comprehensively studying circuit- and network-level function and organization in awake rodents. As more datasets of rodent rsfMRI, potentially collected in separate animal models of brain disorders or under different physiological conditions, become available, these data can be integrated for further investigations of circuit- and network-level changes in these rodent models.

## Acknowledgments

The present study was partially supported by National Institute of Neurological Disorders and Stroke (R01NS085200, PI: Nanyin Zhang, PhD) and National Institute of Mental Health (R01MH098003 and RF1MH114224, PI: Nanyin Zhang, PhD).

## Appendix

## Folder structure of the database

Fig. 9 shows the folder structure of the database. Data of each subject are placed in a single folder named ‘rat’ followed by the subject index ‘xxx’. In each folder, raw data, unprocessed rsfMRI scans, preprocessed scans, and intermediate files from the preprocessing are separately placed in the folders named ‘raw’, ‘rfmri_unprocessed’, ‘rfmri_processed’, and ‘rfmri_intermediate’, respectively. All image files are in the NIfTI (Neuroimaging Informatics Technology Initiative) format. Also, sequence name and acquisition time of each scan, as well as their names in each folder are stored in the file named ‘ratxxx_info.json’ in the JSON (JavaScript Object Notation) format. In the ‘rfmri_intermediate’ folder, despiked, coregistered, and motion-corrected images are provided, as well as motion parameters (in the files ended with ‘_motion.txt’) and averaged WM/CSF signal (in the files ended with ‘_WMCSF_timeseries.txt’) from spatially smoothed image. Scans with less than 90% frames left after despiking were discarded. Results from single-scan ICA cleaning and IC labels are placed in a folder ended with ‘.gift_ica’. Each image file in the ‘rfmri_intermediate’ and ‘rfmri_processed’ folders is accompanied with a JSON file containing the processing steps completed and parameters used.

**Figure 9.**
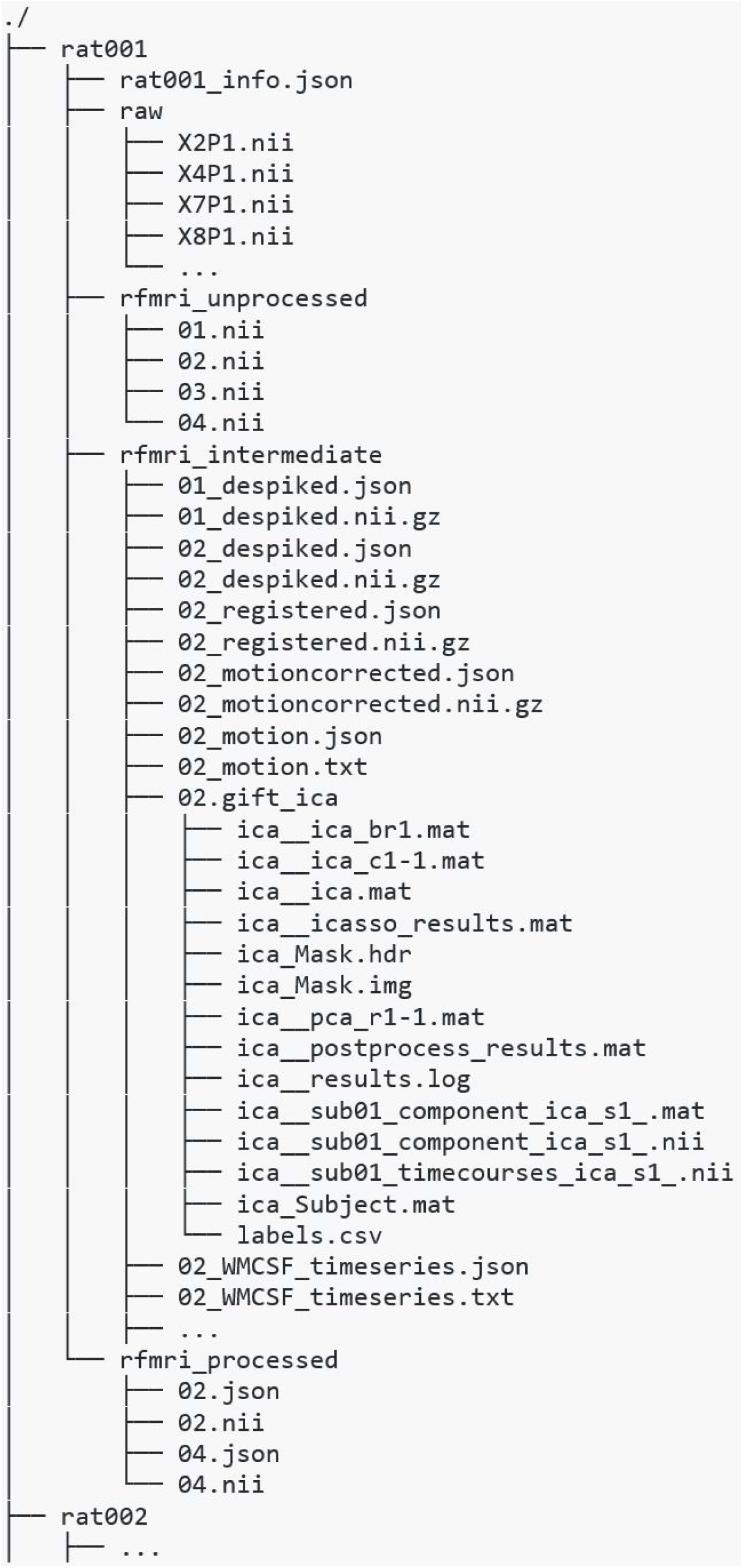
Folder structure of the database.

---

**Figure S1.**
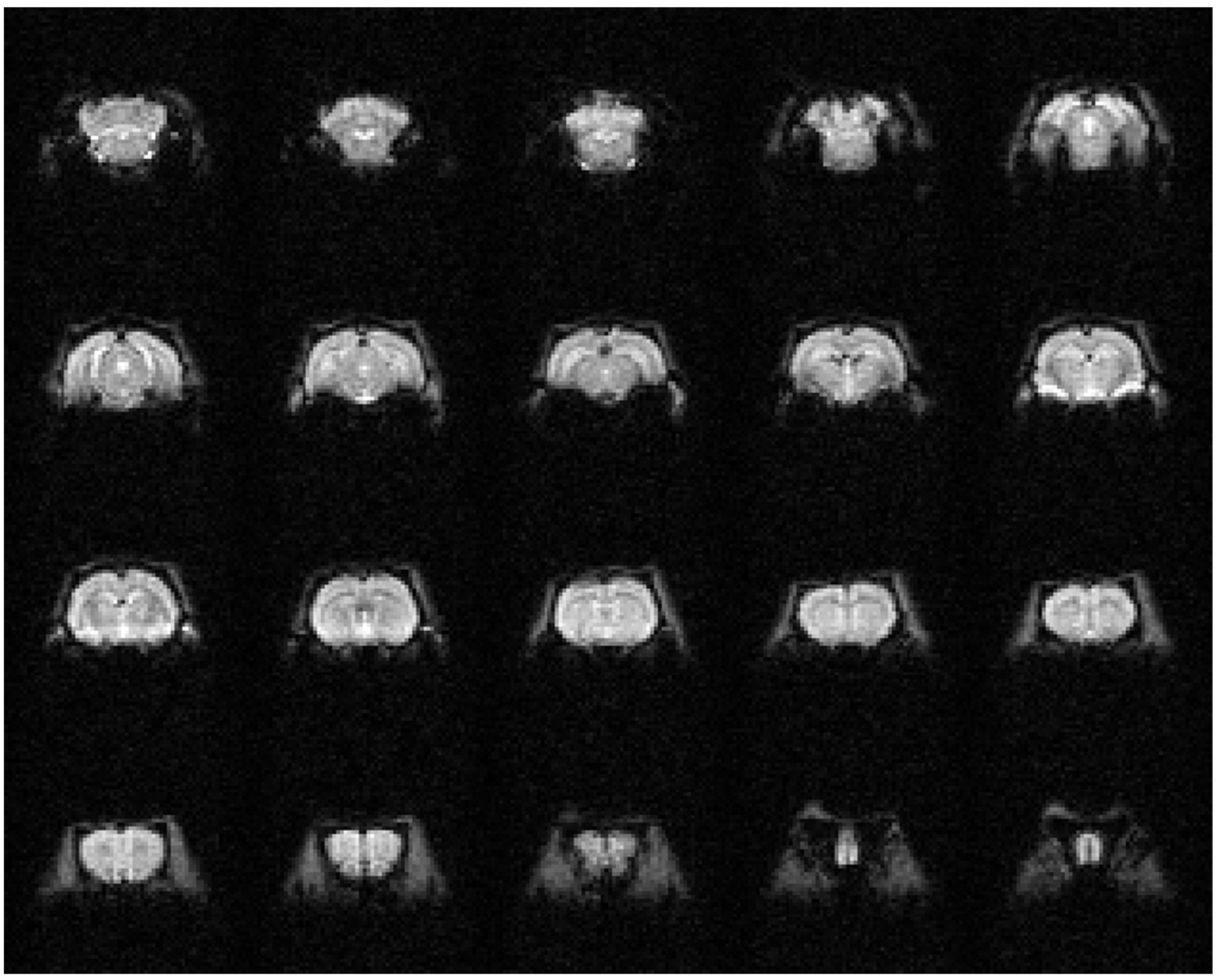
A representative raw rsfMRI frame.

**Figure S2.**
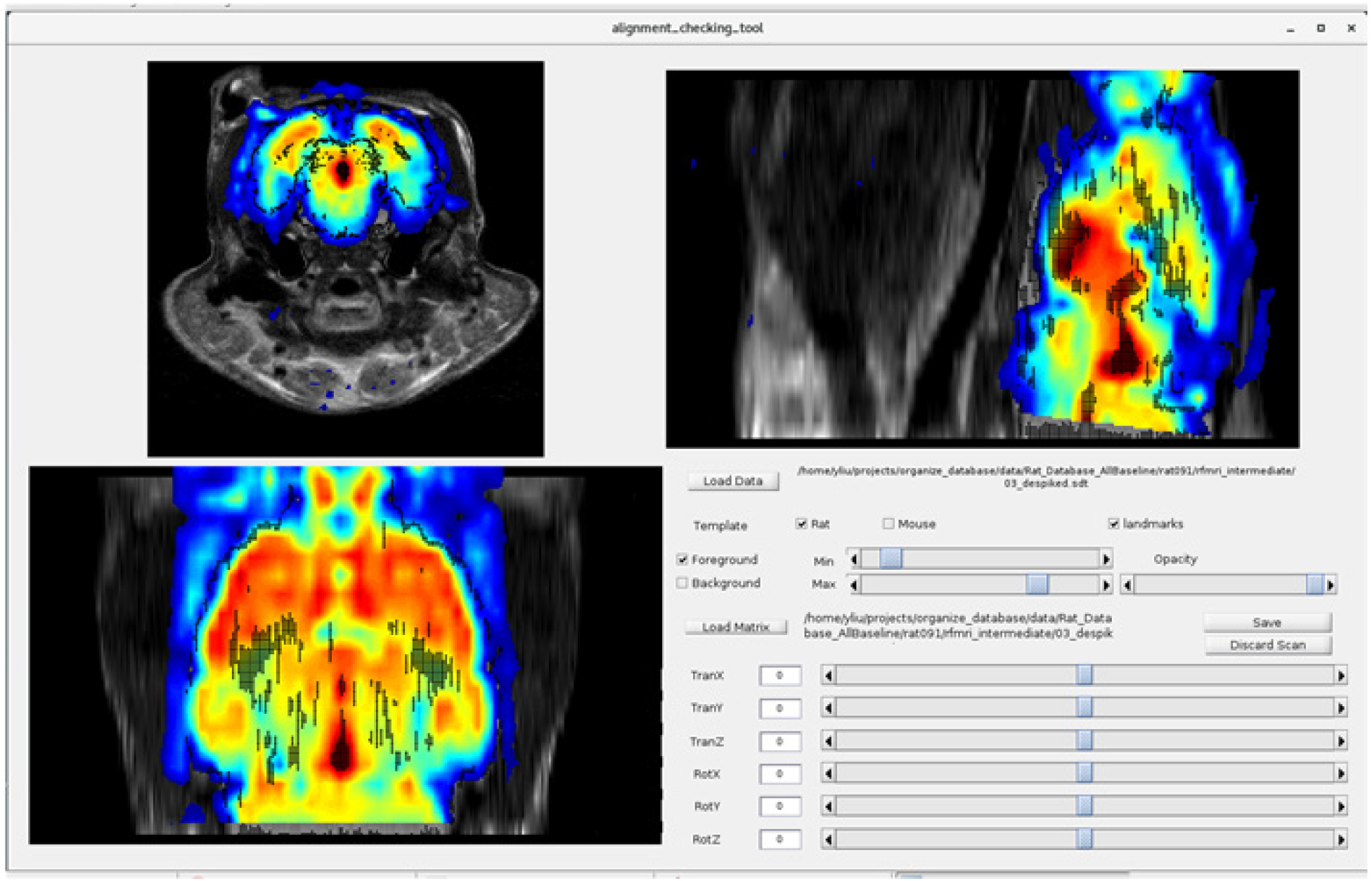
User interface of the coregistration toolbox.

